# Image denoising for fluorescence microscopy by self-supervised transfer learning

**DOI:** 10.1101/2021.02.01.429188

**Authors:** Yina Wang, Henry Pinkard, Emaad Khwaja, Shuqin Zhou, Laura Waller, Bo Huang

## Abstract

When using fluorescent microscopy to study cellular dynamics, trade-offs typically have to be made between light exposure and quality of recorded image to balance phototoxicity and image signal-to-noise ratio. Image denoising is an important tool for retrieving information from dim live cell images. Recently, deep learning based image denoising is becoming the leading method because of its promising denoising performance, achieved by leveraging available prior knowledge about the noise model and samples at hand. We demonstrate that incorporating temporal information in the model can further improve the results. However, the practical application of this method has seen challenges because of the requirement of large, task-specific training datasets. In this work, addressed this challenge by combining self-supervised learning with transfer learning, which eliminated the demand of task-matched training data while maintaining denoising performance. We demonstrate its application in fluorescent imaging of different subcellular structures.

## Introduction

Fluorescence microscopy is an indispensable technique for studying biological dynamics, or detecting and quantifying molecules in subcellular compartments. Recent advances such as light-sheet microscopy [1, 2] and super-resolution microscopy [3] have enabled subcellular fluorescence imaging at a high temporal and/or spatial resolution. Photobleaching of the fluorophores limits the total amount of signal that can be extracted from a biological sample. In addition, for live imaging, phototoxicity, such as failure or delay of cell division or perturbation of biological processes, can occur well before substantial photobleaching is observed [4, 5]. Therefore, a trade-off has to be made between light exposure and quality of the recorded image. In many cases, e.g. when tracking a very rapid process over a long period of time or when the abundance of the target molecule is low, the resultant images could become extremely noisy. In these cases, image denoising is important to enable extracting useful information from the data [6–8].

Deep learning based processing of microscopy images has recently emerged as a powerful approach for a variety of tasks [9–12]. This approach has demonstrated promising denoising performance by learning the structure and noise characteristics for a particular type of sample and experimental condition. Content-Aware Restoration (CARE) is a typical example of this method [8]; a convolutional neural network (CNN) is trained with a large number of noisy and clean image pairs. With this data-driven prior knowledge, the network learns to statistically transform noisy pixels to clean ones. Still, this strategy poses two practical challenges. First, the performance of a deep learning system greatly depends on the amount and quality of the training dataset [6]. Typically, hundreds to thousands of noisy and clean image pairs relevant to the task are needed for fluorescence microscopy denoising. Acquiring such training data sets requires intensive effort and often dedicated experiments [8]. Moreover, it is not always possible to acquire clean images due to intrinsic constraints of certain samples. Second, supervised neural networks often have trouble generalizing to images not present or adequately represented in the training data set. They can easily memorize the training data and be tuned to prior content [6, 8]. However, when applied to real images acquired under different conditions or from other types of cellular structures, hallucination artifacts can occur, producing results that appear real but are incorrect. This issue not only signifies the burden of high quality, matched training data acquisition, but also casts doubt on the ability for denoising by supervised learning to discover previously unknown biological phenomena.

More recently, self-supervised deep learning image denoising methods have been developed to address these challenges [13, 14]. These methods utilize the independence of noise among noisy images of the same sample or pixels across the same image, so that only noisy images are needed for training. This approach eliminates the need to acquire clean training datasets and can even rely solely on the images to be denoised as the training data. However, compared to supervised learning using matching training data sets, the denoising performance of these self-supervised learning methods is degraded due to the absence of prior knowledge that could be learned from the training data set [13].

Here, we first showed that incorporating additional temporal information into supervised learning improves denoising performance, though it still suffers from intrinsic limitations of supervised learning. Then, we present a denoising method that leverages transfer learning to take advantage of both supervised and self-supervised learning. The method first pre-trains a deep neural network with generic and/or synthetic noisy/clean cellular image pairs using supervised learning. It then re-trains this network with the noisy images from the specific task using self-supervised learning. We demonstrate that this scheme can assimilate information regarding image resolution, noise statistics and sample morphology from the two training steps, thus achieving superior denoising performance and robustness than either supervised or self-supervised learning alone, while negating the need for task-specific training datasets.

## Results

### Limited improvement of supervised learning denoising with additional temporal information

In our initial efforts to improve image denoising performance of supervised deep learning, we took advantage of the temporal consistency of structures in living cells. To capture the redundant information across images in a sequence, we used the classical U-Net architecture [15] similar to that in CARE [8] and expanded the network to include time as an additional dimension. We referred to this architecture as timeUnet. (Supplementary Fig. 1. See Supplementary Note for details of the implementation). In timeUnet, a 2D image sequence is treated as a 3D data set, with the network predicting one denoised image from a sliding window of 11 images in the sequence. To benchmark the performance, we generated synthetic noisy image sequences by adding Poisson shot noise and calibrated sCMOS camera noise to experimental high signal-to-noise ratio (SNR) confocal movies of mitochondria in cells. Synthetic images with various SNR, achieved by linearly scaling the image peak intensity to a desired value, were used to test the denoising performance across a range of SNRs. We demonstrate that, rather unsurprisingly, timeUnet out-performed CARE, which denoises individual images (Supplementary Fig. 2). In the entire range of input SNR tested, timeUnet consistently produced lower Mean Averaged Error (MAE) and higher Structural Similarity Index Measurement (SSIM) [16] values when comparing the denoised images to the ground truths (Supplementary Fig. 3). In particular, the addition of time-domain information clearly helped reducing denoising artifacts, correctly recovering images of discrete mitochondria that were connected in the CARE result (Supplementary Fig. 2). Such artifacts could be highly detrimental in the study of certain biological processes such as mitochondria fission and fusion.

Despite the improved performance, timeUnet retains the drawbacks of supervised learning. A large library of paired noisy and clean movies is still needed. Moreover, this library must match the condition of the actual denoising task. For example, any difference between the SNR of the training and test input data leads to a degradation of denoising performance for both CARE and timeUnet (Supplementary Fig. 4a). Particularly, applying the model to test data with higher SNR than the training data actually produced quantitatively worse denoising results than those from test data with matching (and lower) SNR (Supplementary Fig. 4c). This phenomenon is highly concerning for practical applications because it means any variabilities in sample or experimental conditions are detrimental even if they improve input data quality. In contrast, self-supervised learning using noise2self [13] performs more consistently over the SNR range of input data, although in the case when the SNR of training and test data sets are matched, supervised learning clearly yields better results (Supplementary Fig. 4c).

### Image denoising by combining self-supervised learning and transfer learning

In order to combine the benefits of both supervised and self-supervised learning, we connected them using transfer learning. For this purpose, we took advantage of the fact that an identical U-Net architecture can be used for supervised learning [8] as well as self-supervised learning by noise2self [13]. The only difference is that the input and output of supervised training are a pair of noisy and clean images, whereas those of noise2self come from the same noisy image split by a pixel mask. To enable transfer learning, we first trained a network by supervised learning using generic cellular microscopy images. Then, for each denoising task, we retrain this network by noise2self using the task image set (Fig. 1a). We compared the performance to the previous noise2self method, which initializes network parameters with random values for self-supervised training.

**Fig. 1.**
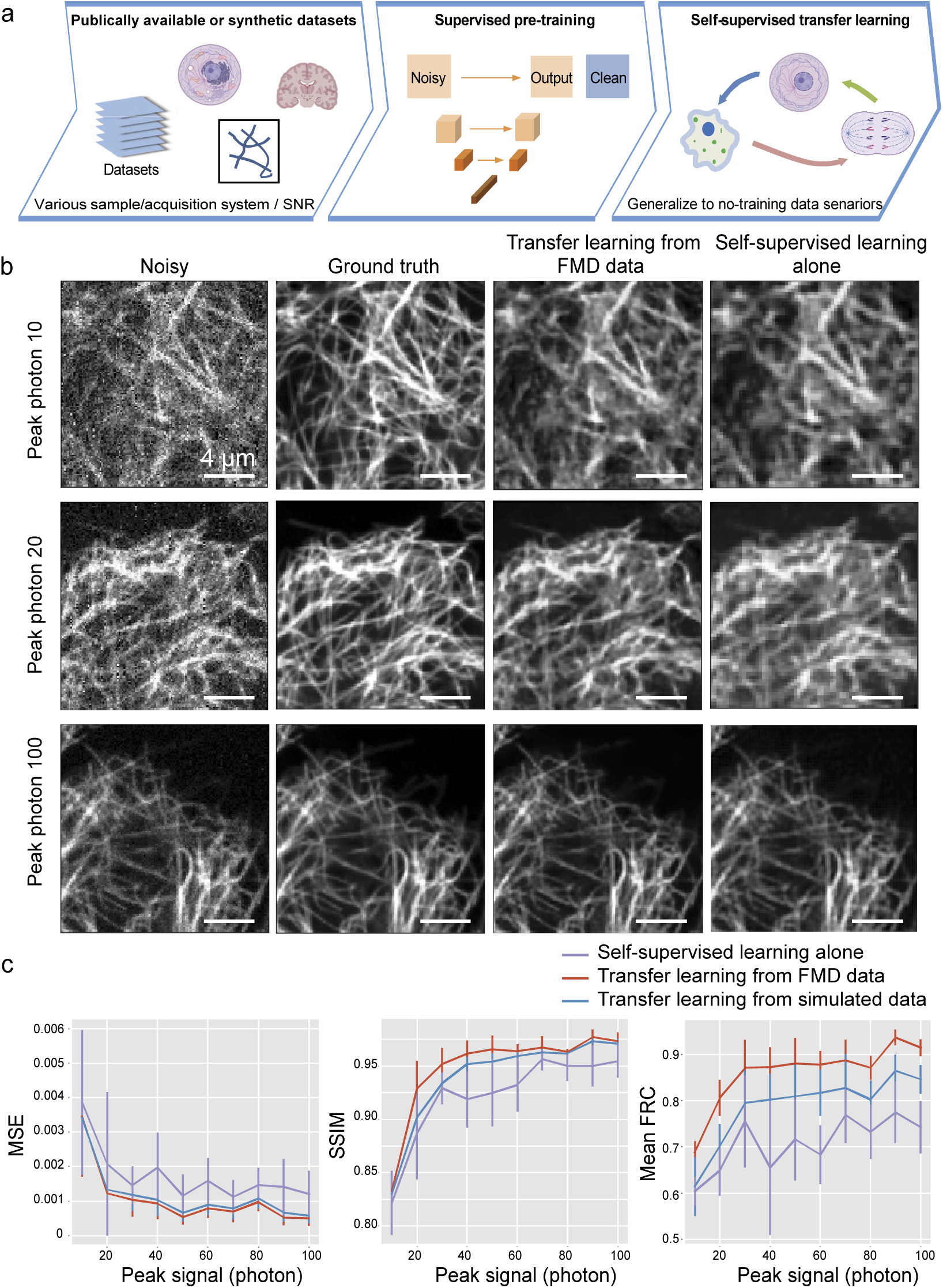
Diagram and performance of transfer learning denoising. (a) Schematic of the transfer learning method. (b) Synthetic noisy images from microtubule confocal images and corresponding denoised images with transfer learning denoising from pre-training using the FMD dataset, compared to self-supervised denoising without pre-training. (c) The denoising performance, in terms of Mean Squared Error (MSE, lower value for less deviation from ground truth), Structure Similarity Index Measurement (SSIM, high value for better similarity with ground truth) and mean Fourier Ring Correlation (FRC, higher value for better resolution of the output image), as a function of the peak signal of synthetic noisy image. Error bars represent standard deviations from 10 test images. The cell and tissue cartoons in (a) were created at BioRender.com.

Specifically, we still used the U-Net architecture. For training at the supervised learning stage, we chose a publicly available dataset, FMD [17], which contains pairs of noisy and clean images from various sample features (subcellular structures of nuclei, F-actin, mitochondria; brain slices and zebrafish) and image acquisition settings (confocal, two-photon, wide-field) (Supplementary Fig. 5). The FMD dataset effectively contains 60,000 noisy image realizations from 240 fields-of-view. For the self-supervised learning stage, we retrained the network by noise2self [13] on a test data set of 10 synthetic noisy images generated from high SNR experimental images in a similar way as described earlier. The retrained network generated denoised images for the same test data set (Fig. 1(b)). Compared to self-supervised learning alone (random initialization of parameters) and using the original high-SNR images as ground truth, transfer learning decreased the Mean Square Error (MSE) loss and provided a higher (Fig.1(c)). This indicates that prior information embedded in the pretrained network contributes to inferring lost information in the noisy measurements. The most prominent visual difference, though, is that denoising results by transfer learning clearly had much better effective spatial resolution (Fig. 1(b)), i.e., more recovery of high spatial frequency components. This visual impression was confirmed by quantifying the average Fourier Ring Correlation (FRC) between the denoised images and the ground truth, which quantifies the effective image resolution [18] (Fig. 1(c)). To further test the tolerance on pretraining data, we generated a set of simulated fluorescence microscopy images containing curve lines resembling microtubules (Supplementary Fig. 6) as the pre-training dataset before self-supervised retraining. Although the resulted denoising performance was slightly worse than FMD-pretrained model, it still out-performed the no-pretraining model in all three metrics (Fig. 1(c)).

We performed similar tests on synthetic images based on confocal images of lysosome structures, which are also absent in the FMD dataset. Our results showed the same trends as those from microtubule images, with transfer learning clearly outperforming self-supervised learning without pretraining in all three metrics (Fig. 2). It is also evident that, in the case of pre-training using simulated images of curved lines, self-supervised retraining on noisy lysosome images can allow the model to correctly restore the punctate appearance of lysosomes despite their drastic morphology difference from the curved lines used in pretraining. Another benefit of pretraining is the stability of performance in repetitive tests on the same set of synthetic noisy lysosome images. For transfer learning self-supervision, each self-supervised training rerun gave almost exactly the same output (indicated by the almost invisible error bar in Fig. 2(b)); meanwhile the output from the no-pretraining model is not stable at all due to the random initialization. Thus, transfer learning also acts as a stabilization of the denoising performance.

**Fig. 2.**
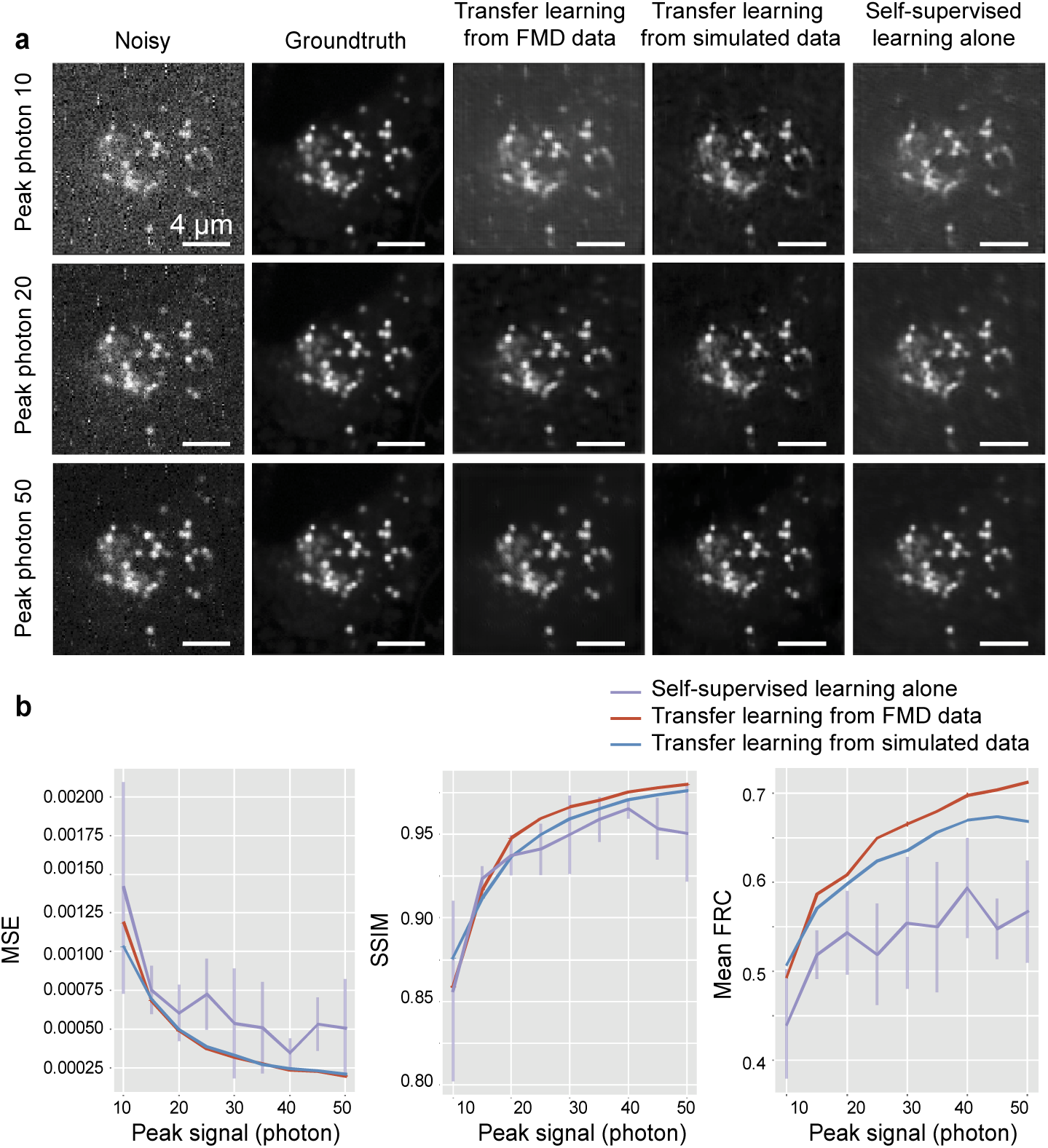
Performance of self-supervised deep denoising with transfer learning on lysosome images. (a) Synthetic noisy images from ground truth lysosome confocal images and denoising results from transfer-learning models pretrained with either the FMD dataset or simulated curved-line dataset, compared to denoising by self-supervised learning along without pre-training. Noisy images with three different levels of peak signal (photons) are shown. (b) The denoising performance, in terms of MSE, SSIM and mean FRC, as a function of the peak signal of the input noisy image. Error bars represent standard deviations from 10 repeated re-runs of the self-supervised training.

To identify what was learned by the model during the pretraining and retraining stages, we applied the FMD- and simulated-pretrained model directly to the denoising of microtubule and lysosome images without self-supervised retraining. The denoised images clearly had numerous morphological artifacts (Supplementary Fig. 7). The FMD-pretrained model output did not display well-defined structures, whereas the simulation-pretrained model generates short segments of curved lines (despite that lysosomes should appear as small spots). Such artifacts are understandable because neither microtubule nor lysosome were present in the images we used from FMD for training, and the simulated pretraining data were curved lines. Compared to the transfer learning results, the retraining using self-supervised learning effectively adapted the pretraining models to the morphology and noise statistics of unseen application data. On the other hand, the major difference between self-supervised denoising with or without transfer learning is the image resolution as measured by FRC. It suggests that self-supervised training implicitly learned a lower image resolution than the actual image resolution because of the corruption of high-frequency information in the images by the noise, while supervised learning on clean images correctly learns the image resolution. This knowledge on image resolution can be effectively transferred to the re-trained model.

### Robustness of transfer learning denoising

To characterize the robustness our approach, we tested the effect of self-supervised training data sizes on denoising performance of microtubule images (Fig. 3). Surprisingly the denoising performance does not show an obvious increase with the increase of training data size in the ranges of 1 to 50 test images, which indicates single-shot self-supervised denoising is possible. We also evaluated the denoising performance when L1 loss is used for the self-supervised training. Generally, L1 loss can better restore sharp features from noisy data compared to L2 loss. In this case, the denoising performance from transfer learning self-supervised model is still much better than the no-pretraining model (Fig. 4), indicating that the source of performance improvement is not in the loss function but because of transfer learning. To appreciate the limits of our proposed training strategy (Fig. 1(a)), we note that our method has limited denoising performance for images with extremely low SNRs. At extremely low SNR conditions, very little useful information is in the image and there is not an adequate prior encoded in the pretrained network parameters. This problem potentially can be solved by including more low-SNR images in the pretraining dataset.

**Fig. 3.**
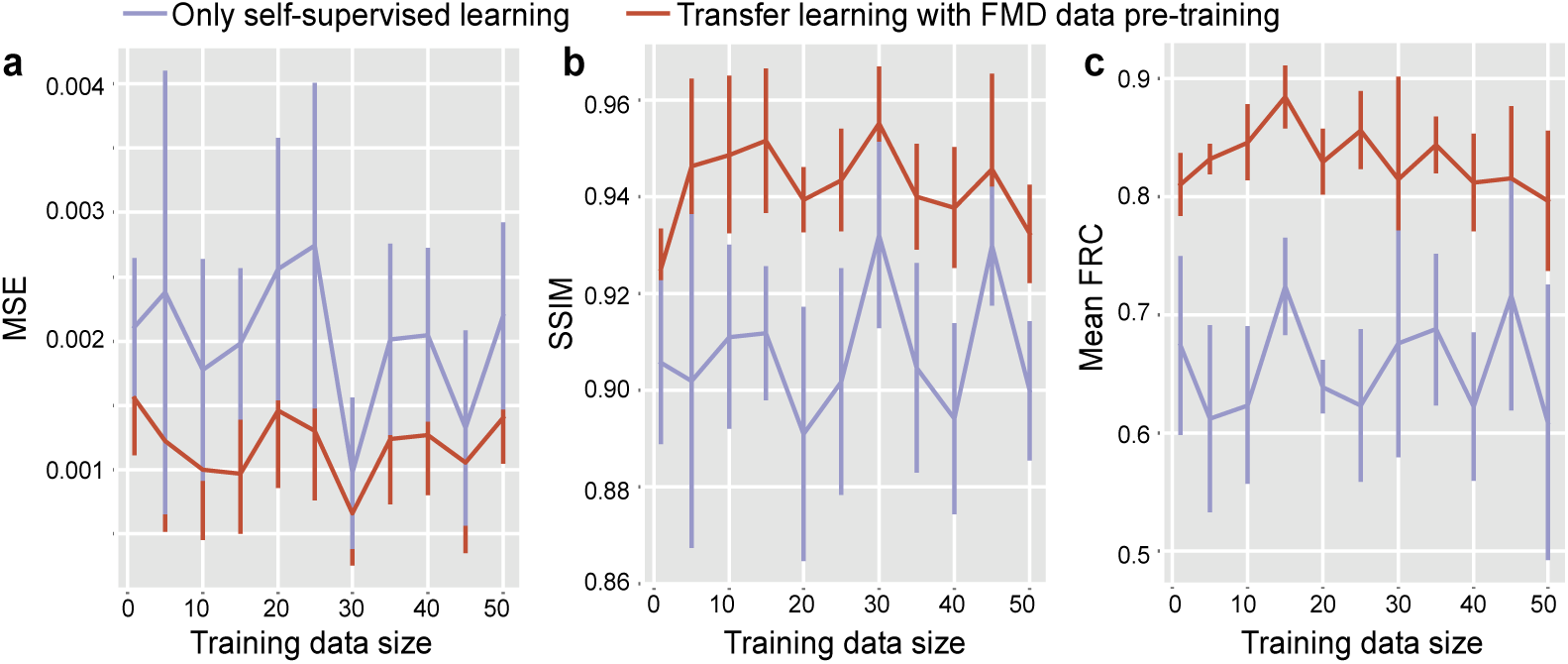
The MSE, SSIM and mean FRC denoising performance as a function of training data size used during the self-supervised training phase. Error bars represent standard deviations from 10 test images.

**Fig. 4.**
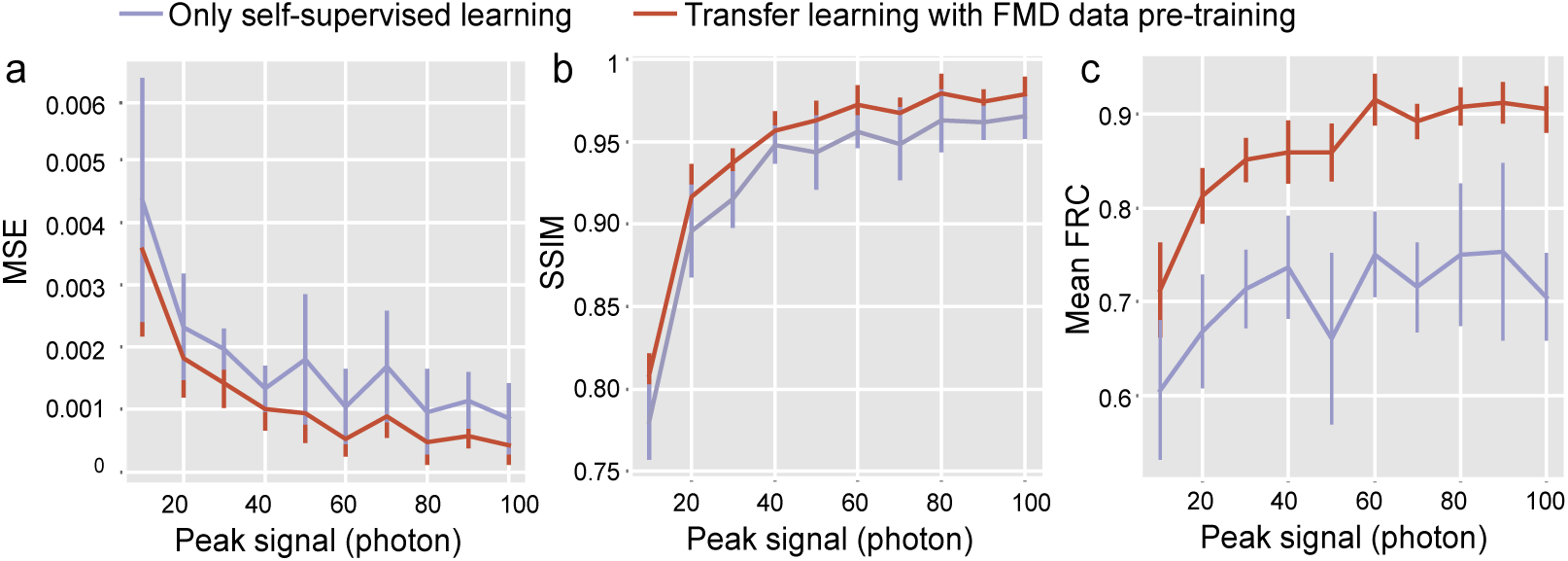
The MSE, SSIM and mean FRC denoising performance as a function of peak signal in synthetic noisy images when L1 is used as loss function instead of L2. Error bars represent standard deviations from 10 test images.

## Discussion

Despite tremendous successes, there are two major barriers for employing deep learning in biological investigations. First, current deep learning approaches typically do not generalize well to data obtained from different samples (Supplementary Fig. 7), different microscopes or under different image acquisition conditions (Supplementary Fig. 4) [6]. While large, diverse training datasets can help solve many of these issues, they are often unavailable for biological studies. On the other hand, the transfer learning approach [11, 19] can practically enable cross-study generalization and information sharing. Second, generating noisy and ground truth image pairs for supervised training is not always practical. For example, for live sample imaging, it is impossible to acquire such image pairs without system hardware modifications. For this reason, self-supervised learning becomes very attractive in eliminating the need for ground truth images. As a result, we took the approach of transfer learning to blend supervised training and self-supervised training to make our method practical and robust. We demonstrated that knowledge of the image high-frequency components are transferrable from the supervised to self-supervised learning phase. With this transferred information, we achieve blind denoising for new tasks just using a few snapshots of noisy images.

## Methods

### Dataset for supervised pretraining

We used Fluorescence Microscopy Dataset (FMD) data set to perform the supervised pretraining. The dataset is downloaded from [17]. We choose this dataset because: (1) the dataset consists of representing images of multiple commonly imaged types of samples (cells, zebrafish, and mouse brain tissues) and multiple commonly used imaging modalities (commercial confocal, two-photon, and wide-field microscopes); (2) the dataset has multiple signal-to-noise ratio realizations of the same imaged scenes. The dataset composes of images from 240 field-of-views (FOV). For each FOV, 50 noisy camera frames were taken, and then image averaging was used to effectively generate the ground truth image and noisy images with various SNR.

In addition to the FMD dataset, we also generated simulation images of curved lines to test the capability of transferring sharp features of our framework. In the simulation, each image is generated by firstly convolving a normalized Gaussian point spread function (standard deviation of 1 pixel) with an object image that consists of random 10 curved lines with various lengths. Then the peak intensity in the image is linearly scaled to a desired value, and Poisson noise and calibrated sCMOS readout and gain noise is added to the image.

### Generating synthetic subcellular structure images for self-supervised training and testing

In order to evaluate the performance of our framework, we first acquired high SNR confocal images of subcellular structures of mitochondria and lysosome. We seeded human HEK 293T cells on an 8-well glass bottom chamber (LabTek). In order to achieve better cell attachment, 8-well chamber was coated with Poly-L-Lysine (Sigma-Aldrich) for 20 mins before seeding cells. For microtubule staining, SiR-tubulin dye (Cytoskeleton) was added directly to the culture medium (100 nM final concentration) and incubate overnight before imaging. For lysosome staining, LysoTracker Blue DND-22 (Thermo Fisher Scientific) was added directly to the culture medium (50 nM final concentration) and incubate for 30 mins before imaging. Live-cell imaging was acquired on an inverted Nikon Ti-E microscope (UCSF Nikon Imaging Center), a Yokogawa CSU-W1 confocal scanner unit, a PlanApo VC 100x/1.4NA oil immersion objective, a stage incubator, an Andor Zyla 4.2 sCMOS camera and MicroManager software.

Then we synthesized noisy images from high SNR clean images to perform the self-supervised training and evaluate the results. The peak intensity in the image is again linearly scaled to a desired peak signal value, and Poisson shot noise and calibrated sCMOS readout and gain noise (based on the camera specifications) was added to the image using a Gaussian random number generator.

### Neural network architecture and training

We used an U-Net architecture implemented in noise2self paper [13, 15]. Each convolutional block consisted of two convolutional layers with 3×3 filters followed by an InstanceNorm. The number of channels was [32, 64, 128, 256]. Down-sampling used strided convolutions and up-sampling used transposed convolutions. The network was implemented in PyTorch. For supervised training, the loss is mean square error. Learning rate was set to 0.001. We trained with a batch size of 32 and 50 epochs.

We used the noise2self self-supervised training strategy. In practice, a masked image, in which a selected subset of pixels was set to zeros, was output to the neural network. The loss was only evaluated on the coordinates of that subset of pixels that are set to zeros. During training, the masked pixels were cycled to make sure a heterogeneous denoising performance over the whole image. In this way, the training process avoids identical map of the input and only relies on the independence between pixels. In this self-supervised training, the loss can be in a different form with supervised training if necessary. The learning rate was 0.0001, an order of magnitude smaller than supervised learning for network parameters fine-tuning. Because the self-supervised training was done use a few snapshots of noisy images alone, these images were processed in a single batch and early stopping with a patience number of 8 used.

### Data and code availability

Python codes for timeUnet, generation of the simulated training data, transfer learning and quantification of model performances, together with test data, are available at https://github.com/BoHuangLab/Transfer-Learning-Denoising/.

## Acknowledgements

We thank Joshua Batson and Loic Royer for inspirational discussions and help with the self-supervised learning code. B.H. is supported by the National Institutes of Health (R01GM131641 and R01GM124334), the UC Berkeley – UCSF Sackler Faculty Exchange program, the UCSF Byers Award in Basic Science. L.W and B.H. are Chan Zuckerberg Biohub Investigators.

## References

1. Chen, B.-C., et al., Lattice light-sheet microscopy: Imaging molecules to embryos at high spatiotemporal resolution. Science, 2014. 346(6208): p. 1257998.

2. Yang, B., et al., Epi-illumination SPIM for volumetric imaging with high spatial-temporal resolution. Nature Methods, 2019. 16(6): p. 501–504.

3. Gwosch, K.C., et al., MINFLUX nanoscopy delivers 3D multicolor nanometer resolution in cells. Nature Methods, 2020. 17(2): p. 217–224.

4. Icha, J., et al., Phototoxicity in live fluorescence microscopy, and how to avoid it. BioEssays, 2017. 39(8): p. 1700003.

5. Laissue, P.P., et al., Assessing phototoxicity in live fluorescence imaging. Nature Methods, 2017. 14(7): p. 657–661.

6. Belthangady, C. and L.A. Royer, Applications, promises, and pitfalls of deep learning for fluorescence image reconstruction. Nature Methods, 2019. 16(12): p. 1215–1225.

7. Carlton, P.M., et al., Fast live simultaneous multiwavelength four-dimensional optical microscopy. Proceedings of the National Academy of Sciences, 2010. 107(37): p. 16016–16022.

8. Weigert, M., et al., Content-aware image restoration: pushing the limits of fluorescence microscopy. Nature Methods, 2018. 15(12): p. 1090–1097.

9. Christiansen, E.M., et al., In Silico Labeling: Predicting Fluorescent Labels in Unlabeled Images. Cell, 2018. 173(3): p. 792–803.e19.

10. Nehme, E., et al., Deep-STORM: super-resolution single-molecule microscopy by deep learning. Optica, 2018. 5(4): p. 458–464.

11. Wang, H., et al., Deep learning enables cross-modality super-resolution in fluorescence microscopy. Nature Methods, 2019. 16(1): p. 103–110.

12. Wu, Y., et al., Three-dimensional virtual refocusing of fluorescence microscopy images using deep learning. Nature Methods, 2019. 16(12): p. 1323–1331.

13. Batson, J. and L. Royer, Noise2Self: blind denoising by self-supervision. Preprint at https://arxiv.org/abs/1901.11365, 2019.

14. Krull, A., T.O. Buchholz, and F. Jug, Noise2Void—learning denoising from single noisy images. Preprint at https://arxiv.org/abs/1811.10980, 2018.

15. Ronneberger, O., P. Fischer, and T. Brox. U-Net: Convolutional Networks for Biomedical Image Segmentation. 2015. Cham: Springer International Publishing.

16. Zhou, W., et al., Image quality assessment: from error visibility to structural similarity. IEEE Transactions on Image Processing, 2004. 13(4): p. 600–612.

17. Yide, Z., et al., A Poisson-Gaussian Denoising Dataset with Real Fluorescence Microscopy Images. Preprint at https://arxiv.org/abs/1812.10366, 2018.

18. Koho, S., et al., Fourier ring correlation simplifies image restoration in fluorescence microscopy. Nat Commun, 2019. 10(1): p. 3103.

19. Wang, J., et al., Data denoising with transfer learning in single-cell transcriptomics. Nature Methods, 2019. 16(9): p. 875–878.

