## Supplementary Notes and Figures for "Image denoising for fluorescence microscopy by self-supervised transfer learning"

#### Add temporal information to supervised deep learning denoising

Deep learning image reconstruction in microscopy typically does not use temporal information. The predictions are made frame by frame independently, and thus successive reconstructed images are not necessarily temporally consistent. Meanwhile, supervised deep learning image reconstruction is potentially subject to hallucination artifacts. These artifacts become clearly noticeable when examined across time. We found that adding the temporal dimension (Fig. S1) in the training dataset would lead to more accurate reconstructions. Specifically, we build an architecture called timeUnet as depicted by Fig. S1. The input to the network is a short video (we used video of 11 frames); the spatial and temporal information of this video is encoded by sets of 3D convolution; then the encoded 3D features are flattened by convolution and the flattened images are used for transpose convolution decoding. Thus, the output of the network is a single reconstructed image targeting a clean image corresponding to the middle frame of the short video. We implemented timeUnet in Keras. We noted that similar idea was used in literature [1] to improve the performance of image super-resolution.

To test the performance of timeUnet in reducing reconstruction artifacts, we used synthetic mitochondria dataset. Both training and testing datasets are generated from mitochondria confocal images with the same procedure descripted in the main text. Firstly, the timeUnet network is trained with a training dataset of 30 videos (batch size: 32; learning rate: 0.001; epoch: 100). Then it was tested on a different set of synthetic mitochondria images. As a result, timeUnet is able to reduce reconstruction artifacts (Figs. S2-S4) and delivers systematically lower MSE error and higher SSIM across a range of SNR conditions compared with the CARE method [2] in which no temporal information is used. Though adding temporal information enforces temporal consistency in image reconstruction, the strategy requires a bigger amount of training data as well as noisy and clean image pairs for the same dynamic.

### Supplementary figures

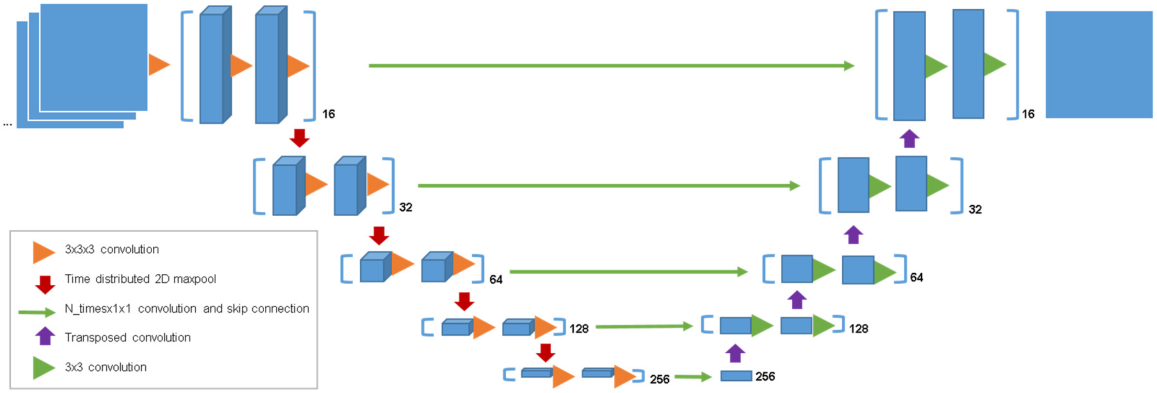


**Fig. S1** Schematic of the timeUnet architecture, with the input being a stack of 11 images in a time sequence and the output being one denoised image at the middle time point.

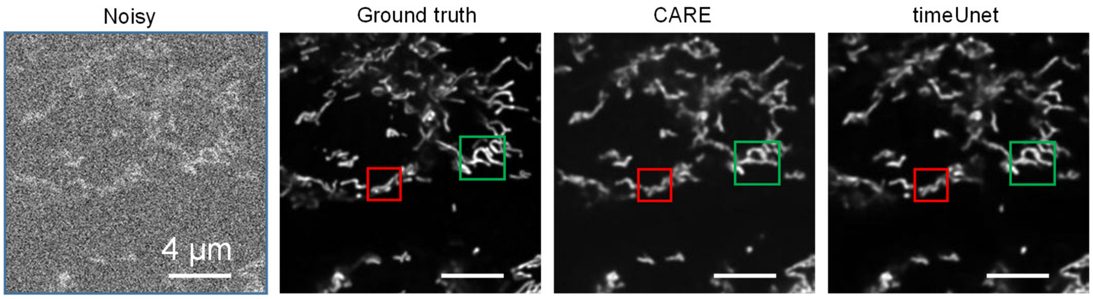


**Fig. S2** Synthetic noisy image from mitochondria confocal images is shown. The peak signal in photon of the noisy image is 10 with simulated Poisson and calibrated camera noise added. Corresponding denoised images from CARE and timeUnet are shown correspondingly. The boxed regions show timeUnet results in smaller number of artifacts than CARE when compared with the ground truth.

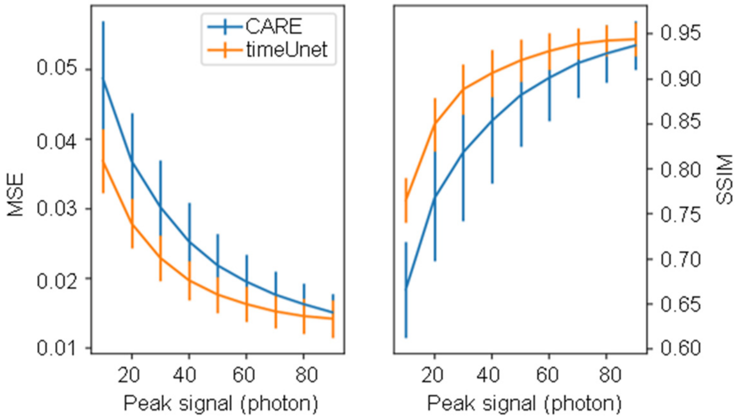


**Fig. S3** The denoising performance, in terms of mean square error, structural similarity index measurement, as a function of the peak signal of synthetic mitochondria noisy images. Error bars represent standard deviations from 10 test data.

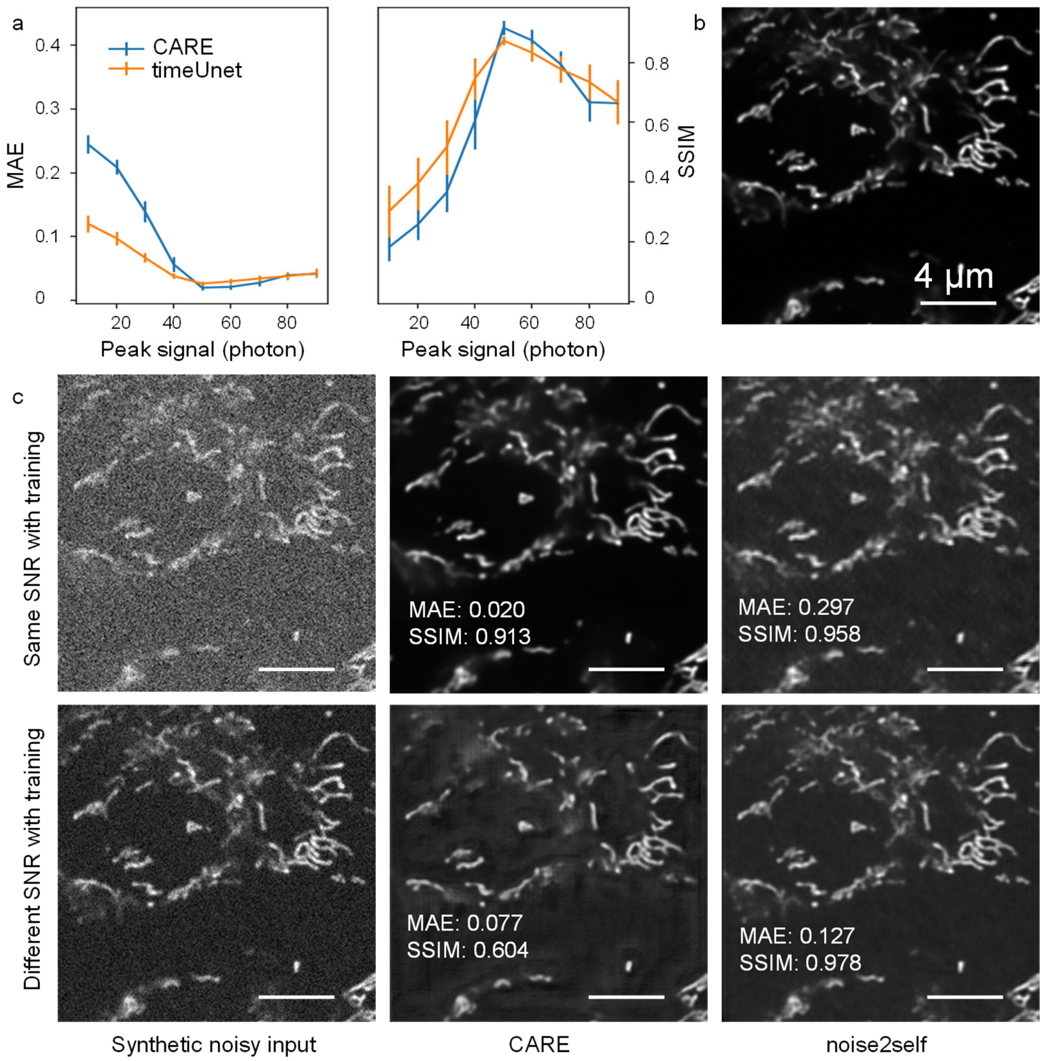


**Fig. S4** Generalization problem of supervised learning image denoising. (a) The performance, in terms of mean absolute error and structural similarity index measurement, of the CARE and timeUnet methods on various peak signal levels, given the networks were trained with training dataset only consists of synthetic images with peak signal of 50 photons. The training dataset was synthetically generated from randomly cropping ~5k 256 × 256 image patched from 30 2048 × 2048 confocal movies. The results show that there is a clear performance peak at test condition that exactly the same with training condition and therefore demonstrating the generalization problem of supervised image denoising. (b) An example cropped region from ground truth confocal images. (c) The upper row displays synthetic test image with the training SNR condition, the CARE method successfully denoised the image and outperformed self-supervised denoising method noise2self. The bottom row displays synthetic test image with SNR better than training condition (peak signal 80 photons). Theoretically the bottom row noisy image is easier to denoise. However, the CARE method provided poorer denoising performance in this case, while noise2self exhibits more stable denoising performance. Error bars represent standard deviations from 10 test data.

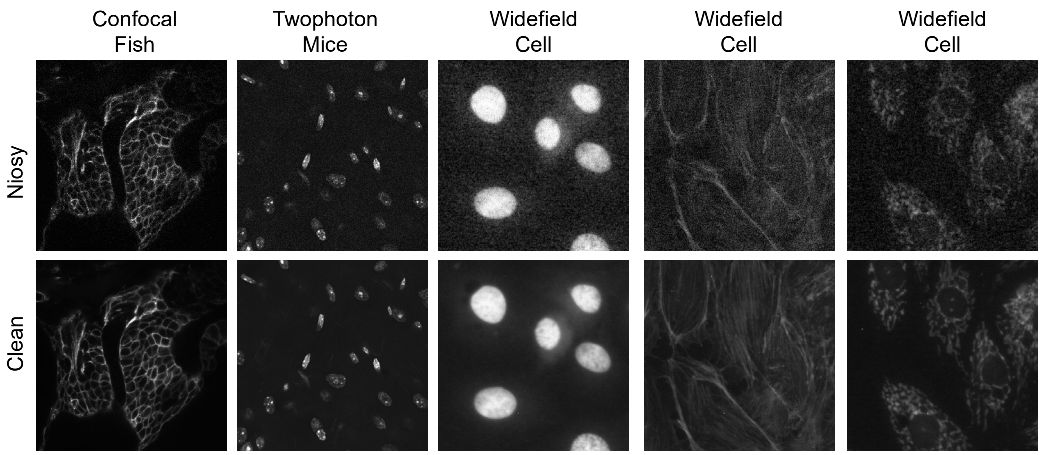


**Fig. S5** Example images of the FMD dataset, including confocal image of zebrafish embryo, two-photon image of nucleus in mice brain tissue, widefield images of nuclei, F-actin and mitochondria of BPAE cells.

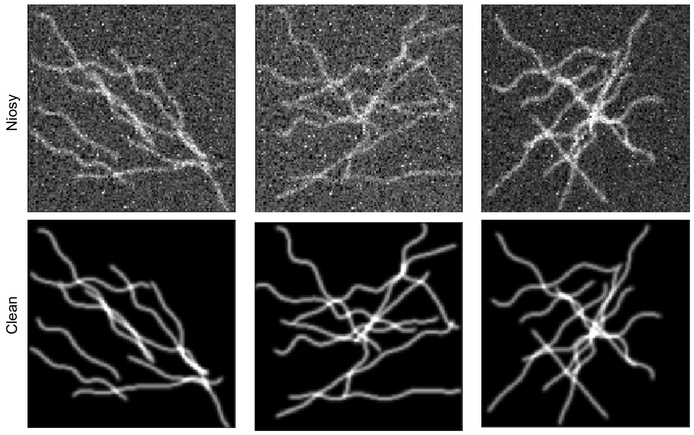


**Fig. S6** Example images consist of curvy lines of the simulation dataset.

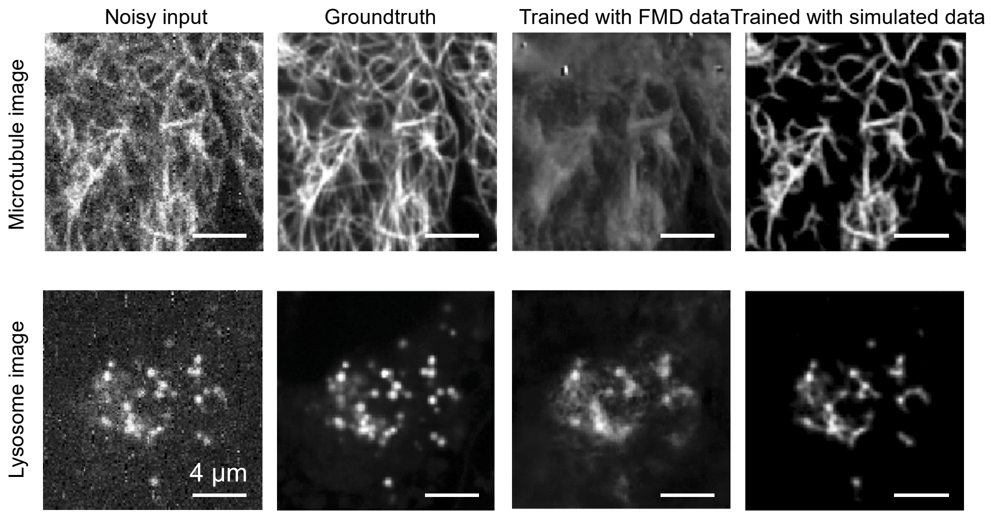


**Fig. S7** Denoising with pretrained networks using FMD or simulated dataset.

**References**

1. L. Fang, F. Monroe, S. W. Novak, L. Kirk, C. Schiavon, S. B. Yu, T. Zhang, M. Wu, K. Kastner, Y. Kubota, Z. Zhang, G. Pekkurnaz, J. Mendenhall, K. Harris, J. Howard, and U. Manor, "Deep Learning-Based Point-Scanning Super-Resolution Imaging," bioRxiv, 740548 (2019).
2. M. Weigert, U. Schmidt, T. Boothe, A. Müller, A. Dibrov, A. Jain, B. Wilhelm, D. Schmidt, C. Broaddus, S. Culley, M. Rocha-Martins, F. Segovia-Miranda, C. Norden, R. Henriques, M. Zerial, M. Solimena, J. Rink, P. Tomancak, L. Royer, F. Jug, and E. W. Myers, "Content-aware image restoration: pushing the limits of fluorescence microscopy," *Nature Methods* **15** (12), 1090-1097 (2018).
